# Loss of *TP73* function contributes to amyotrophic lateral sclerosis pathogenesis

**DOI:** 10.1101/451419

**Authors:** Jonathan M. Downie, Summer B. Gibson, Spyridoula Tsetsou, Kristi L. Russell, Matthew D. Keefe, Karla P. Figueroa, Mark B. Bromberg, L. Charles Murtaugh, Joshua L. Bonkowsky, Stefan M. Pulst, Lynn B. Jorde

**Author notes:** Correspondence to: 15 S 2030 E RM 5100, Salt Lake City, UT 84112, USA. These authors contributed equally to the manuscript (co-first author). Co-senior author.

## Abstract

Much remains unknown about the genetics and pathophysiology underlying the neurodegenerative disease amyotrophic lateral sclerosis (ALS). We analyzed exome sequences from a cohort of 87 sporadic ALS (SALS) patients and 324 healthy individuals. *TP73*, a homolog of the *TP53* tumor suppressor gene, had five rare deleterious protein-coding variants; in a separate collection of >2,900 ALS patients we identified an additional 19 rare deleterious variants in *TP73*. An *in vitro* C2C12 myoblast growth assay confirmed that these variants impair or alter *TP73* function. *In vivo* mutagenesis of zebrafish *tp73* using CRISPR led to impaired motor neuron development and abnormal axonal morphology, concordant with ALS pathology. Together, these results demonstrate that *TP73* is a risk factor for ALS, and identifies a novel dysfunctional cellular process in the pathogenesis of ALS.

## Main Text

Amyotrophic lateral sclerosis (ALS) is a fatal degenerative disease of motor neurons in the brain and spinal cord^1^. Much remains unknown about the genetics and pathophysiology underlying this disease. Genetic factors are critical determinants of ALS pathogenesis, as 68% of familial ALS (FALS) and 17% of sporadic ALS (SALS) cases have an identifiable genetic risk variant^2; 3^. However, up to 61% of SALS risk has been attributed to genetic factors^4; 5^, suggesting that unknown genetic loci contribute to the development of SALS. To address this, we employed next-generation sequencing of a cohort of 87 SALS patients, together with *in vitro* and *in vivo* experiments, to identify and characterize the role of the *TP73* gene, a member of the p53 family of transcription factors, in ALS. This discovery reveals an unexpected and previously undescribed dysfunctional cellular process involved in ALS pathogenesis.

To identify novel candidate ALS risk loci, we obtained whole exome sequences from a cohort of 87 SALS patients from the University of Utah and 324 healthy control individuals from the Simons Simplex Collection^3; 6^. *VAAST*^7^ was used to identify genes with excess deleterious nonsynonymous single nucleotide variants (SNVs) in ALS patients compared to unaffected controls. We excluded SNVs with an Exome Aggregation Consortium (ExAC)^8^ non-Finnish European (NFE) minor allele frequency (MAF) above 0.001. *PHEVOR*^9^, a tool designed to identify disease-relevant candidate genes in small sequencing studies, was then used to re-rank the *VAAST* candidate genes on the basis of biochemical, cellular, and pathological functions similar to known ALS risk genes. Two known ALS risk genes, *MAPT* (rank: 3) and *SOD1* (rank: 5), were among the top ten genes ranked by combined *VAAST* and *PHEVOR* analysis (Figure 1).

**Figure 1.**
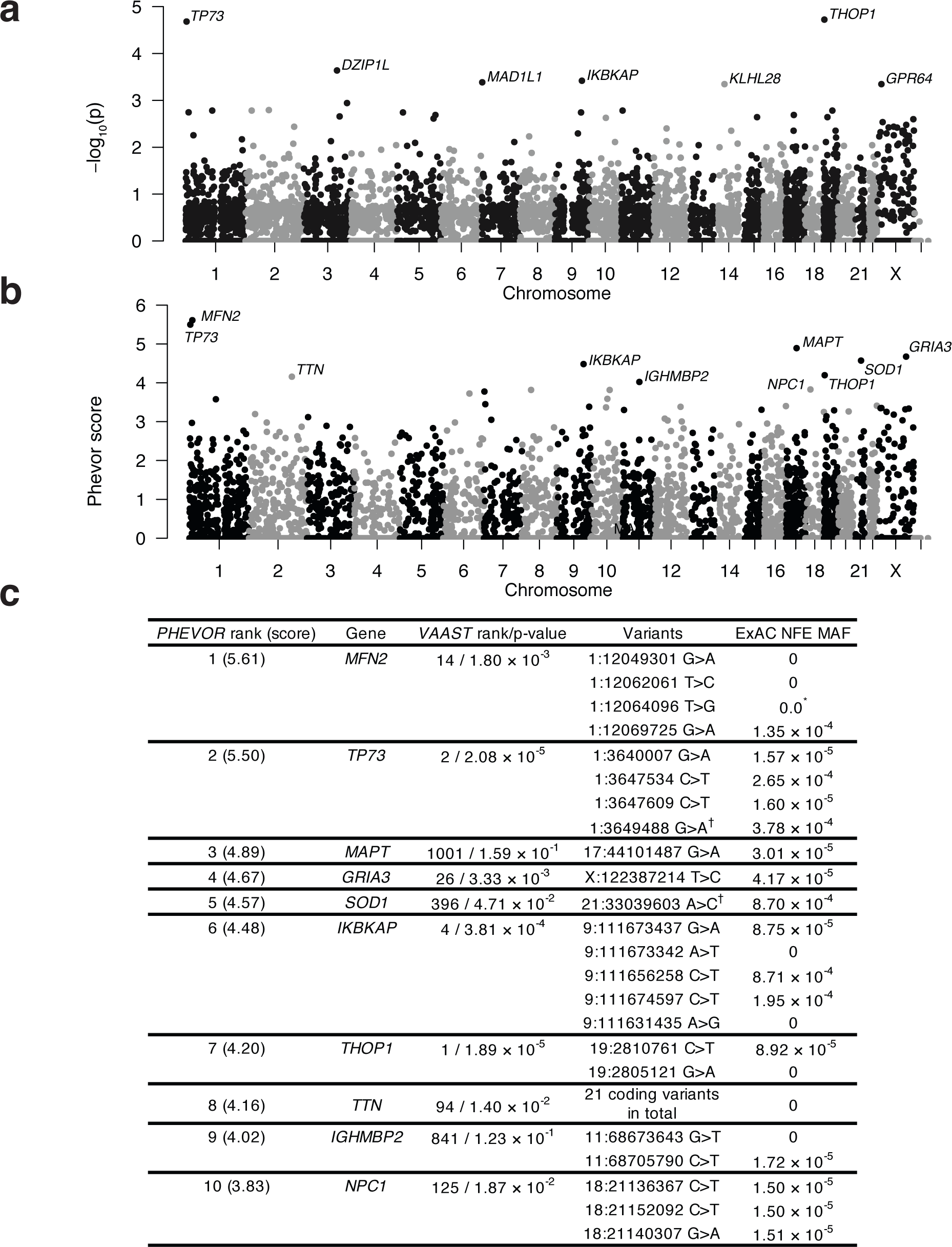
*VAAST*/*PHEVOR* analyses identify multiple genes burdened by deleterious variation in ALS patients. a) Manhattan plot of *VAAST* analysis. Two genes, *TP73* and *THOP1*, approached genome-wide significance (p = 2.57 × 10^-6^ (Bonferroni correction)) b) Manhattan plot of *VAAST*/*PHEVOR* analysis. The *PHEVOR* score is the log_10_ ratio comparing the likelihood a gene is associated with disease vs. healthy individuals. c) Top ten ranked genes from the *VAAST*/*PHEVOR* analysis. The SNVs that are contributing genetic burden are listed (GRCh37). * indicates a variant found populations outside of NFE. †variant found in two SALS patients.

The only gene to rank among the top five genes in both *VAAST* and *PHEVOR* was *TP73* (Figure 1), which encodes tumor protein 73 (p73). *TP73*’s *VAAST* score approached genome-wide significance after Bonferroni multiple test correction, and *PHEVOR* revealed that *TP73* shares significant functional characteristics with known ALS genes. Four different rare missense SNVs in *TP73* were found among five SALS patients (Figure 1c). In contrast, only three rare missense SNVs in *TP73* were found in the Simons Simplex Collection control cohort. An additional rare-seven amino acid in-frame deletion variant (chr1:3646605 CCATGAACAAGGTGCACGGGGG>C; *TP73*:p.PMNKVHGG413-420P), which was not assessed by *VAAST*, was found in an SALS patient upon searching for indels. Five of the six *TP73* protein-coding variants were confirmed by Sanger sequencing; one variant was an exome genotyping error (Supplementary Figure 1).

p73 is part of the p53 family of tumor suppressor transcription factors, which modulate the expression of target genes to affect cell-cycle arrest, apoptosis, and cellular differentiation^10; 11^. *Trp73*^-/-^ mice exhibit brain defects without a spontaneous tumor phenotype^12^, demonstrating that p73 is required for central nervous system development^11^. Aged mice carrying one *Trp73* null allele (*Trp73*^+/-^) show progressive motor weakness and a reduced number of neurons in the motor cortex^13^.

We next tested whether rare and deleterious *TP73* protein-coding variants could be found in two replication cohorts. A *TP73* missense SNV (chr1:3647559 G>A; *TP73*:p.A472T) was identified and verified by Sanger sequencing (Supplementary Figure 1) in an independent cohort of 53 whole-exome or whole-genome sequenced University of Utah ALS patients (Figure 2). An additional 18 rare variants that alter the *TP73* coding sequence were found in the ALSdb cohort (ALSdb, http://alsdb.org) of more than 2,800 whole-exome sequenced ALS patients^14^. Combining all analyzed cohorts, 24 different rare *TP73* coding sequence variant sites (22 SNVs and 2 in-frame indels) were found among approximately 2,900 ALS patients (Figure 2), which is similar to the relative contribution of many other ALS risk genes^2^. All 22 SNVs were predicted to be deleterious to *TP73* function by MetaSVM^15^ and had an NFE MAF < 0.0005 (Figure 2). Furthermore, 14 of these 22 SNVs were considered to be deleterious by CADD^16^ (CADD score ≥ 20). In comparison, 66 of the 242 rare coding SNVs found in ExAC were considered to be deleterious by CADD, which is a significantly lower proportion than the variants identified in the ~2,900 ALS patient cohort (Fisher exact test *p* < 0.001).

**Figure 2.**
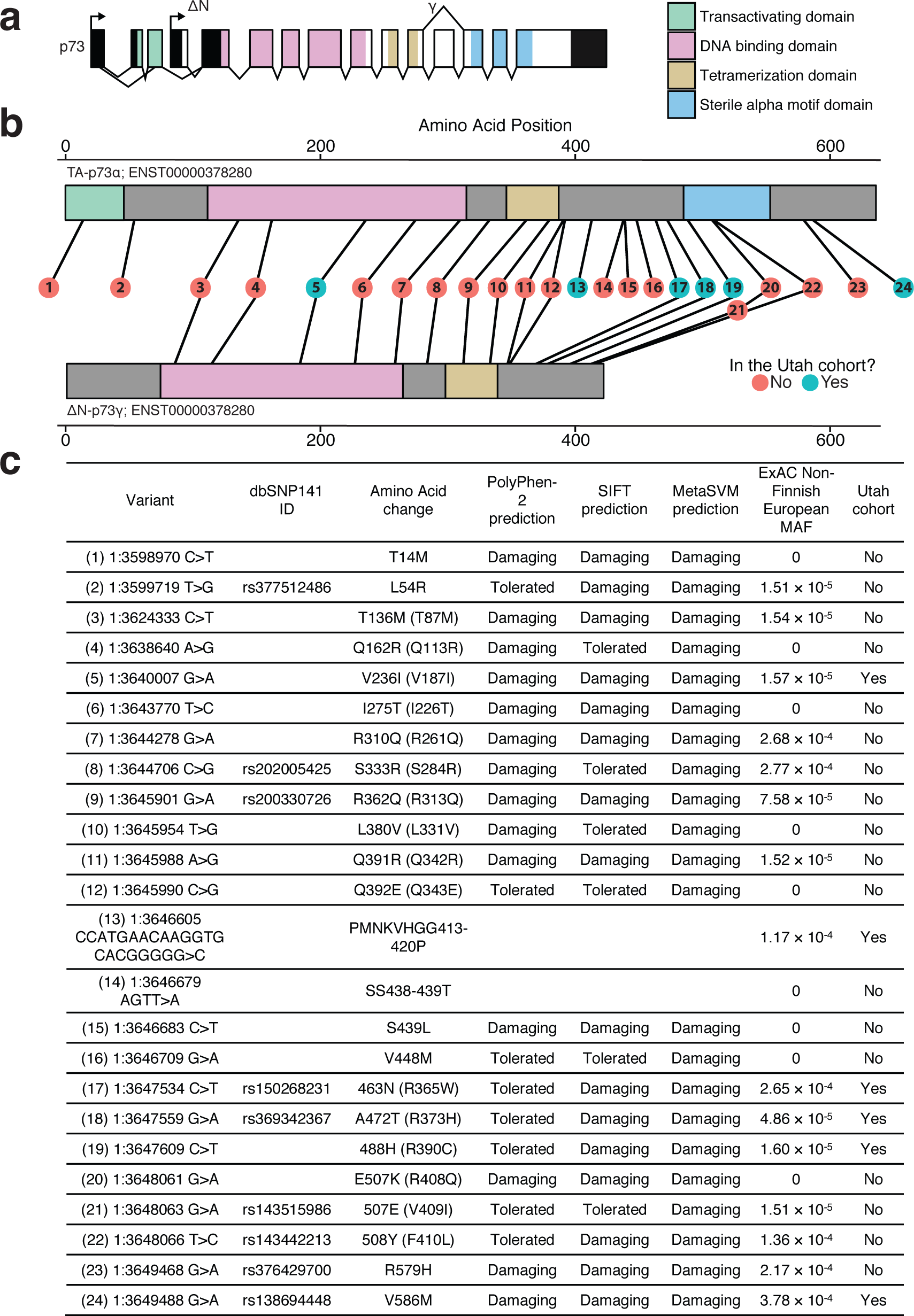
24 rare (ExAC NFE MAF < 0.001) *TP73* amino acid alerting-variants were found across all studied ALS cohorts. a) TA-p73 and ΔN-p73, which have opposing tumor suppressive/oncogenic functions, are expressed from two separate promoters. Alternative splicing events also give rise to a multitude of different isoforms (such as ΔN-p73α and ΔN-p73γ). Adapted from Moll and Slade 2014. b) The primary structure of TA-p73α and ΔN-p73γ. The number within each circle refers to the variant number in c. Four of the nonsynonymous SNVs are not found in p73α proteins, but do exist in p73γ due to splicing differences. c) Information of all 24 *TP73*coding variants. Amino acid positions are listed for TA-p73α (ΔN-p73γ); ENST00000378295 (ENST00000378280).

Next, we tested whether the identified *TP73* protein-coding variants impair the function of p73. *TP73* gives rise to two different main protein isoforms—TA-p73 and ΔN-p73—which have opposing functions^11^. TA-p73, which possesses an N-terminal transactivation domain, induces expression of gene targets^11^. Conversely, N-terminally truncated p73 (ΔN-p73) has oncogenic and transformative qualities because it inhibits the function of p53 and TA-p73^11^. Of note, ΔN-p73 is the primary p73 isoform found in the brain and promotes neuronal survival by providing resistance to apoptotic insults^17; 18^. ΔN-p73 function can be assayed by overexpression in serum-deprived C2C12 myoblasts as it inhibits cell cycle withdrawal, impairs myoblast differentiation into myotubes, and decreases myosin heavy chain (MHC) staining^19^. All four *TP73* variants identified in University of Utah ALS patients that also resulted in coding changes in ΔN-p73α, the canonical isoform ΔN-p73, were selected for functional testing. We tested the ability of wild-type (WT) ΔN-p73α and each selected variant to inhibit myoblast differentiation and MHC staining. Expression of WT ΔN-p73 in C2C12 cells resulted in impaired differentiation, consistent with previous studies^19^ (Figure 3). Cells expressing two of the four tested *TP73* variants did not show inhibited differentiation, suggesting these mutations render ΔN-p73α nonfunctional (Figure 3). Three of the four tested *TP73* variants resulted in a significant increase in myotube width (Figure 3), which suggests hyperfusion of developing C2C12 cells and disordered maturation. Thus, these *TP73* protein-coding variants impair or alter the function of p73.

**Figure 3.**
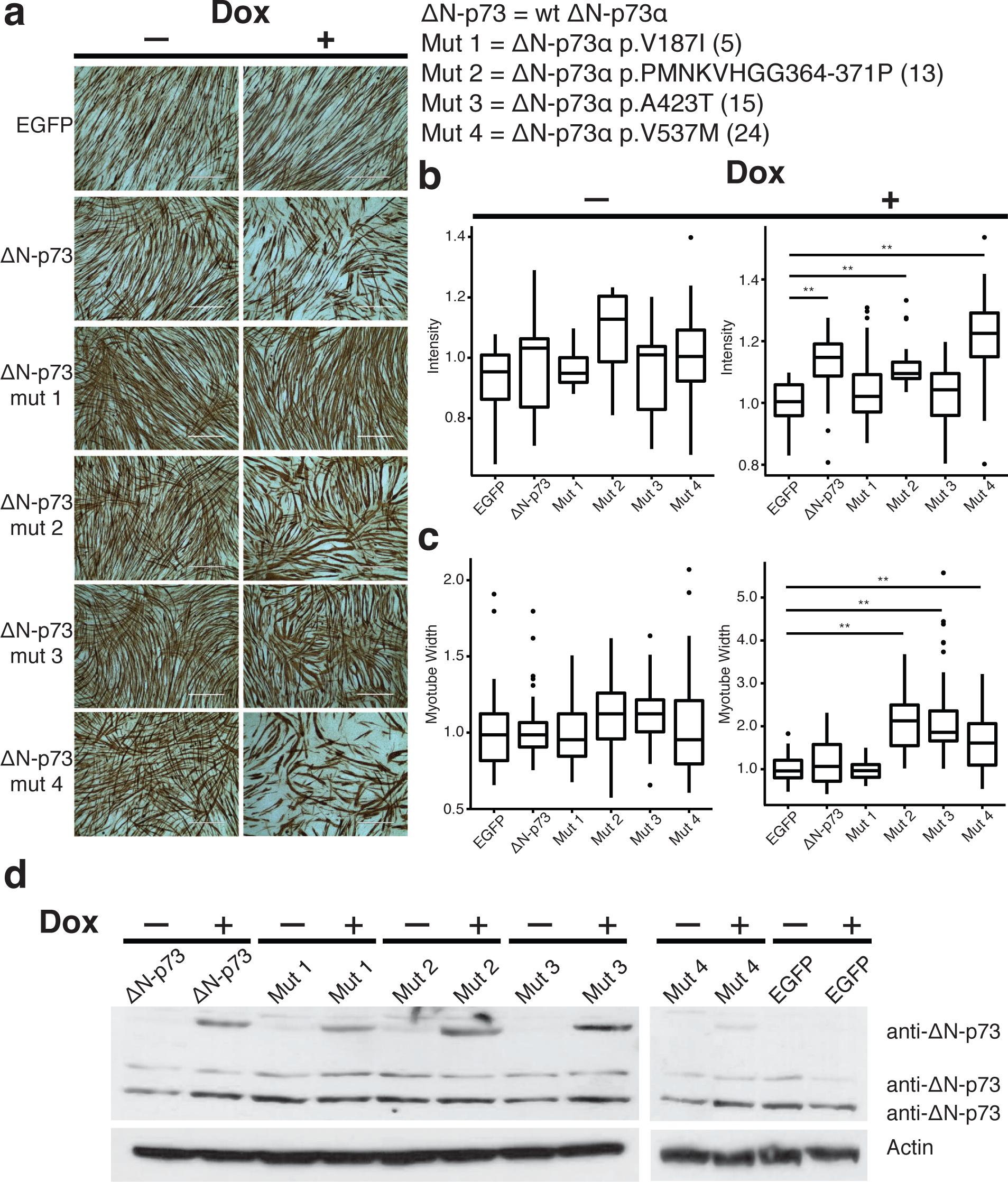
*TP73* ALS variants alter or impair p73 function. a) Doxycyline (Dox) induced WT-ΔN-p73α expression results in impaired C1C12 differentiation shown by reduced MHC IHC staining compared to EGFP. ΔN-p73α^Mut1+3^ expression fails to inhibit C1C12 differentiation shown by no difference in MHC staining compared to EGFP. ΔN-p73α^Mut2-4^ expression results in myotubes with significantly larger width and disordered differentiation compared to EGFP. Amino acid positions are listed according to ENST00000378288. Numbers in parentheses refer to the variant numbers in Figure 2c b) Quantification of normalized white light intensity (inversely related to MHC staining) **=*P*<0.001 (Wilcoxon rank-sum test with Bonferroni correction) c) Quantification of normalized myotube width. d) Western blot showing that ΔN-p73α expression is induced in C2C12 transduced cells when exposed to doxycycline. Additional ΔN-p73 rows likely represent other C-terminal isoforms expressed from germline.

Next, we tested the effect of loss of p73 function on motor neuron development and morphology. Previous studies have established zebrafish (*Danio rerio*) as an ALS model^20^. A CRISPR/Cas system was used to determine whether loss of *tp73* in zebrafish leads to an ALS-like phenotype. Using a guide RNA (gRNA) against exon 4 of zebrafish *tp73* (*tp73*^CRISPR^), which encodes a portion of the DNA binding domain, we found that CRISPR-injected embryos had >95% deleterious mutations at the target locus (Figure 4c). To determine whether mutagenesis of *tp73* disrupts motor neuron development, we quantified GFP^+^ spinal motor neurons (SMNs) in Tg(*Hb9*:*Gal4*; *UAS*:*GFP*) embryos. As a control, we used a gRNA targeting tyrosinase (*tyr*^CRISPR^)^21^. *tp73*^CRISPR^ mutagenized embryos had a significant reduction in the number of GFP-labeled SMN compared to uninjected controls or *tyr*^CRISPR^ injected controls (Figure 4g-j). These results demonstrate the impact of deleterious *tp73* mutations on SMN development, illustrating a pathogenic correlation with motor neuron disease.

**Figure 4.**
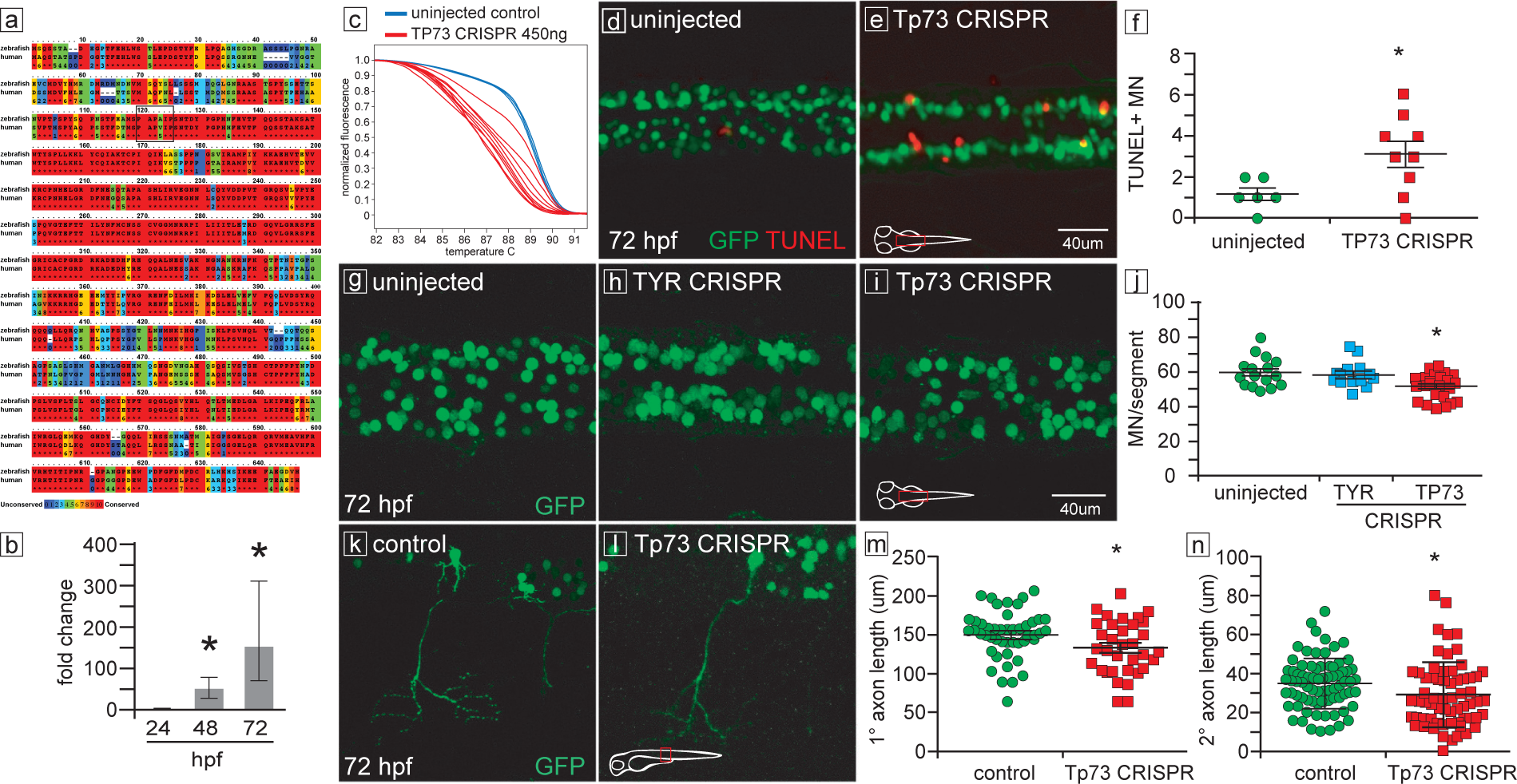
*tp73* mutation impairs motor neuron development and axon outgrowth. a) Conservation of zebrafish and human p73 proteins. Color indicates degree of amino acid conservation (red = conserved, blue = non-conserved). b) qRT-PCR of *tp73*gene expression during development. c) *tp73*CRISPR targeting of exon 4 produced >95% mutagenesis, shown by HRMA genotyping at 24 hpf. d, e) Increased motor neuron (green) apoptosis (red TUNEL labeling) in *tp73*^CRISPR^ embryos compared to uninjected, performed in *mnx:GFP* transgenic line. Confocal imaging, dorsal views, spinal cord. f) Quantification of motor neuron apoptosis in d, e (Uninjected 1.2 +/- 0.3 (Number of animals (n)=6), *tp73* CRISPR 3.1 +/- 0.6 (n=9), *P*<0.05). g-i) *tp73*^CRISPR^ injected embryos have a reduced number of motor neurons compared to both uninjected and *tyr*^CRISPR^ sibling controls, j) Quantification of g-i (Uninjected 59.9 +/- 2.1 (Number of animals (n)=16), *tyr*^CRISPR^ 58.7 +/-2.0 (n=14), *tp73*^CRISPR^ 52.1 +/- 1.5 (n=24), *P*<0.01). k, l) Motor neuron primary axon length in *tp73*^CRISPR^ embryos is significantly shorter than uninjected sibling controls. Motor neurons and their processes are mosaically-labeled with *mnx:GFP*. Confocal images, lateral views, spinal cord. m, n) Quantification of motor neuron primary and secondary axon lengths in *tp73*^CRISPR^ compared to uninjected sibling controls (Primary axon lengths: Uninjected 149.4 +/- 4.4 µm (Number of animals (n)=47), *tp73*^CRISPR^ 132.6 +/- 5.9 µm (n=34), *P*<0.05; Secondary axon lengths: Uninjected 34.7 +/- 1.5 µm (n=76), *tp73*^CRISPR^ 29.0 +/- 2.0 µm (n=64), *P*<0.05).

Previous studies have shown that p73 is required to prevent neuronal apoptosis^18; 22^. To determine whether the reduction in motor neuron number in *tp73^CRISPR^* mutagenized animals was due to apoptosis, we quantified TUNEL^+^/GFP^+^ co-labeled SMNs in Tg(*Hb9*:*Gal4*; *UAS*:*GFP*) embryos. We found a ~2-fold increase in TUNEL^+^ GFP^+^ SMN in *tp73^CRISPR^* mutagenized animals compared to uninjected controls (Figure 4d-f). This suggests that increased apoptosis in *tp73*^CRISPR^ mutants contributes to motor neuron number reduction.

Several ALS zebrafish models^20; 23-25^ have reported a significant reduction in SMN axon length early in development. To address SMN axon outgrowth, we injected *mnx:GFP* alone or in combination with *tp73*^CRISPR^ to quantify SMN axon length. We found that *tp73*^CRISPR^ mutagenized embryos exhibited a significant reduction in primary and secondary axonal branching length (Figure 4k-n). Taken together, these data indicate disruption of *tp73* leads to a reduction in SMN number and axon branch length in development.

Our results show that deleterious *TP73* protein-coding variants occur in ALS patients at a similar frequency to other known risk genes, that *TP73* ALS variants impair normal function of p73, and that loss of p73 impairs motor neuron survival and axonal development. Our data indicate that *TP73* is a novel ALS risk gene. Transcription factors that drive neuronal cell survival, differentiation, and tumor suppressor pathways have not been previously implicated in ALS. These findings reveal unexpected aspects of the ALS genetic risk and pathology landscape and may open new approaches toward treatment.

## Acknowledgments

We thank the Utah Genome Project for providing funds and resources to support next-generation sequencing of our ALS cohort. We would like to give special recognition to the Utah Neuroscience Initiative for funding the *in vitro* and *in vivo* experiments performed in this study. We would like to thank the University of Utah Mutation Generation and Detection Core for the help they provided in generated the CRISPR construct used in the experiments outlined in this manuscript. The work performed in this study was also supported by the following grants from the National Institutes of Health (NIH): TL1TR001066, R01GM059290, R01GM104390, R35GM118335, and R37NS033123. JLB was supported by the Bray Chair in Child Neurology Research and R21MH107039. The content is solely the responsibility of the authors and does not necessarily represent the official views of the NIH. Partial funding support was also provided for S.M.P. by Target ALS.

The control data used in this analysis can be found in the National Database for Autism Research (https://ndar.nih.gov/) under study DOI:10.15154/1149697.

## Author Contributions

Dr. Downie and Dr. Gibson has made a substantive contribution to the design and conceptualization of the study, analysis and interpretation of the data, and drafting of the manuscript. Dr. Tsetsou has made a substantive contribution to the analysis/interpretation of the *in vivo* zebrafish experiments and drafting of the manuscript. Ms. Russell has made a substantive contribution to the analysis/interpretation of the *in vitro* myoblast experiments and drafting of the manuscript. Dr. Keefe has made a substantive contribution to the analysis/interpretation of the *in vivo* zebrafish experiments. Mrs. Figueroa has made a substantive contribution to the analysis and interpretation of the data. Dr. Bromberg has made a substantive contribution to the revising of the manuscript and enrollment of patients into this study. Dr. Murtaugh, Dr. Bonkowsky, Dr. Pulst, and Dr. Jorde has made a substantive contribution to the design and conceptualization of the study, interpretation of the data, and revising of the manuscript.

## Methods

### Samples and sequencing

The exome sequencing results of 87 European sporadic ALS (SALS) patients and 324 SSC control individuals from a previous study were used for this analysis^3; 6^. The detailed sequencing methods and quality-control measures can be found in that report. Experiments were approved by the University of Utah Institutional Review board. SALS patients seen at the University of Utah School of Medicine provided consent and were included in genetic studies. All patients were determined to have probable or definite ALS according to El Escorial revised criteria^26^. Exome sequencing libraries were generated from patient DNA using the Gentra Puregene Blood Kit (Qiagen) and SeqCap EZ Exome Enrichment Kit (v3.0; Roche NimbleGen). In total, DNA from 96 SALS patients was gathered for genetic analysis. The patient characteristics, ancestry composition, and known genetic risk factors—including disease-associated *ATXN2* and *C9orf72* repeat expansions—of this ALS cohort have been previously described^3^. Genomic reads were also obtained from 714 healthy individuals from the Simons Simplex Collection (SSC) to serve as controls^6^.

Exome sequencing libraries were sequenced on the Illumina HiSeq platform to generate 101-bp, paired-end reads. BWA-MEM (v0.7.12) was used to perform read alignment to the GRCh37 reference genome^27^. Genomic read coordinate-based sorted and duplicate read marking was performed by Picard Tools (v1.130). A standard GATK best-practices pipeline was then used to generate joint variant calls for each analyzed individual as previously described^3^.

### *VAAST* and *PHEVOR* analysis

The variant calls from 87 European SALS patients and 324 SSC control individuals were then selected for *VAAST*/*PHEVOR* analysis. *ADMIXTURE*^28^ and *smartpca*^29; 30^ were previously used^3^ to determine that these individuals were of European descent and were specifically selected to control for population stratification effects. Genomic regions covered by less than five sequencing reads on average in the SALS and SSC cohorts were omitted from further analysis to control for coverage differences between the two cohorts. Variants with an ExAC^8^ European MAF greater than 0.001 were removed from further consideration to reduce false-positive gene associations. This value corresponds approximately to the frequency of the most common allele known to cause ALS^31^.

Genotype calls passing these criteria were analyzed by *VAAST*^7^ to prioritize genes more burdened by deleterious variants in cases versus controls. Insertion and deletion (indel) variants were not used in the *VAAST* analysis as indel frequency-based filtering is difficult due to inconsistency in how they are reported in different datasets^32^. *VAAST* was run assuming a dominant model of disease inheritance. Multiple test correction is required when using *VAAST* because it tests for an excess of burden in all genes in the genome. Bonferroni correction was used to account for multiple hypothesis testing. As a result, a p-value of 2.57 × 10^-6^ was required for a gene to be considered significantly burdened (*α* = 0.05; 19,492 genes tested). The output from *VAAST* is a ranked list of genes by burden significance. This output is then analyzed by *PHEVOR*^9^ in order to identify burdened genes that share similar functional characteristics to genes known to be associated with a phenotype of interest. *PHEVOR* first collects genes previously shown to be associated with a phenotype as provided by the Human Phenotype Ontology (HPO). *PHEVOR* then traverses multiple gene ontologies—such as the Gene Ontology, Mammalian Phenotype Ontology, and the Disease Ontology—using genes from the HPO gene list to find ontology nodes, and the genes contained within them, likely to be associated with the phenotype. *PHEVOR* then combines the degree of association a gene has with a particular phenotype with the gene burden calculations of VAAST to determine the *PHEVOR* score. The *PHEVOR* score is defined as the log_10_ ratio of how likely a gene is to be associated with a disease phenotype of interest versus a healthy phenotype.

The following Human Phenotype Ontology (HPO)^33^ terms were used by the *PHEVOR* analysis: amyotrophic lateral sclerosis (HP:0007354), abnormal motor neuron morphology (HP:0002450), motor neuron atrophy (HP:0007373), and frontotemporal dementia (HP:0002145).

### Screening of ALS patients for deleterious *TP73* variants and Sanger sequencing

The discovery SALS cohort was screened for indels in *TP73* after performing *VAAST* as these variants were not assessed in this analysis, which found chr1:3646605 CCATGAACAAGGTGCACGGGGG>C; *TP73*:p.PMNKVHGG413-420P in one individual.

We utilized two replication cohorts to determine whether rare and deleterious *TP73* protein-coding variants could be found outside of the discovery cohort. The first was a cohort from the University of Utah that contained exome and whole genome sequenced SALS patients. More specifically, it contained nine exome sequenced non-European SALS patients who underwent genotype calling with the discovery cohort. It also included 44 SALS patients whose whole genomes were sequenced. These individuals came from a University of Utah ALS sequencing project that contained 70 ALS patients and eight unaffected relatives. Of these 70 ALS patients, 26 were also exome sequenced either in the discovery or University of Utah replication cohort and were not considered in this analysis. The collected DNA samples were Illumina whole-genome sequenced to an average coverage of 60X with 150-bp paired-end reads by NantOmics as part of the Utah Heritage 1K Project. Genomic reads from each sequenced individual were then aligned to the GRCh37 reference genome using the BWA-MEM aligner^27^. The aligned genomic reads from all individuals underwent joint variant calling with 95 long-lived individuals (longevity cohort) and 291 European individuals (CEU (Utah Residents (CEPH) with Northern and Western European Ancestry) and GBR (British in England and Scotland)) from the 1000 Genomes Project^34^ using the Genome Analysis Toolkit (GATK; v.3.4-46) best practices guidelines^35-37^. The variant calls from all 53 ALS patients from the University of Utah replication cohort were screened for rare *TP73* coding variants. One additional *TP73* coding variant was found from this cohort (chr1:3647559 G>A; *TP73*:p.A472T).

Sanger sequencing was used to validate the seven identified *TP73* coding variants in the University of Utah ALS patients. Sanger sequencing confirmed the presence of all but one of the *TP73* coding sequence variants. More specifically, the presence of a rare *TP73* SNV (chr1:3649488 G>A; *TP73*:p.V586M) was confirmed in only one of the two ALS patients it was first identified in (Supplementary Figure 1). The primers used for Sanger verification of these variants can be found in Supplementary Table 1.

### Cloning of wildtype and mutant ΔN-p73

A DNA insert containing the ΔN-p73α coding DNA sequence (CDS) was then generated from a HA-p73α-pcDNA plasmid (a gift from William Kaelin; Addgene plasmid # 22102^38^) by PCR (Supplementary Table 1). The coding sequence of ΔN-p73 was cloned into a pBABE-puro vector (a gift from Hartmut Land & Jay Morgenstern & Bob Weinberg, Addgene plasmid # 1764^39^) using BamHI and SalI (New England Biolabs). Patient variants were introduced into pBABE-ΔN-p73-puro using site directed mutagenesis (New England Biolabs) (Supplementary Table 1). Wildtype and mutant ΔN-p73 were amplified and cloned into the Gateway donor vector pDONR221 through BP reactions (Invitrogen). Next, LR reactions between the pDONOR221 vectors and the lentiviral Destination vector pCLX-pTF-R1-DEST-R2-EBR65 (a gift from Patrick Salmon, Addgene plasmid # 45952^40^) were performed. HEK293T cells were transfected using a calcium phosphate protocol with plasmids psPAX2 (a gift from Didier Trono, Addgene plasmid # 12260), pCAG-Eco (a gift from Arthur Nienhuis & Patrick Salmon, Addgene plasmid # 35617) and wildtype or mutant pCLX-pTF-ΔN-p73 added to produce lentivirus. pCLX-pTF-eGFP was used to produce control lentivirus. HEK293T cells were incubated in high glucose Dulbecco’s Modified Eagle’s Medium (DMEM) supplemented with 10% heat inactivated fetal bovine serum (HI FBS, Genesee Scientific), and 1% pen/strep (P/S, Thermo Fisher). Vectors were added to HEK293T cells in growth medium plus 25uM chloroquine. After 24 hours, chloroquine media was removed and replaced with growth medium. Supernatant containing lentivirus was harvested 48 hours post-chloroquine removal.

### C2C12 in vitro testing of ALS *TP73* variants

C2C12 myoblasts were cultured in high glucose DMEM plus 20% HI FBS and 1% P/S. For lentiviral infections, 50,000 C2C12s were seeded per well. Twenty µl of lentiviral supernatant was added to cells in growth medium containing 5 µg/ml polybrene. After 24 hours, medium was replaced with growth medium lacking polybrene. Forty-eight hours post infection, cells were incubated in growth medium plus 10 µg/mL blasticidin to select for successfully infected cells. Cells were continuously kept under blasticidin selection in all subsequent experiments. Twenty-four well plates were prepared for C2C12 differentiation experiments by adding 0.1% gelatin in dH20 to each well and incubating for fifteen minutes at 37C. Wells were rinsed with PBS and wells were seeded with 50,000 C2C12 cells in technical duplicate for each distinct genotype. Cells exposed to doxycycline (dox) were cultured in separate culture dishes from controls. Cells were exposed to either growth medium containing 1 µg/ml of doxycycline or normal growth medium. After 24 hours, cells were incubated in differentiation medium (DM) (10 µg/ml recombinant human insulin (Gibco), 2% HI horse serum (Thermo Fisher), and 1% P/S) with or without 1 µg/ml doxycycline. Cells were incubated in DM with or without doxycycline for four days before fixation with 70% ethanol. DM was changed every 48 hours. After fixation and endogenous peroxidase inactivation, cells were blocked for one hour with blocking solution (5% donkey serum in PBS, 0.3% TX-100, 0.2% Na Azide). Cells were incubated with anti-myosin heavy chain antibody (MF-20, DSHB) at a 1:100 dilution at 4C overnight. Cells were washed with PBST and incubated with donkey-anti mouse conjugated to biotin secondary for 45 minutes at room temperature (Jackson ImmunoResearch). Cells were washed and Vector ABC reagent (Vector Laboratories, VECTASTAIN Elite ABC-HRP Kit #PK-6100) was added to cells and incubated for thirty minutes at room temperature. Cells were washed and Vector DAB substrate was added (Vector Laboratories, DAB Peroxidase (HRP) Substrate Kit #SK-4100). Substrate was allowed to develop for 5 minutes at room temperature. Cells were imaged using an EVOS Plate Reader (imaging was performed at the Fluorescence Microscopy Core Facility, a part of the Health Sciences Cores at the University of Utah). Control plates with no dox exposure were analyzed in biological duplicate and plates with dox exposure were analyzed in biological triplicate. Four images were taken of each technical replicate well for each genotype on both positive and negative doxycycline plates (though some images could not be used to measure either intensity or myotube width due to technical inadequacy). Three distinct measurements of myotube width were performed per image. Myotube width was measured using pixel length in Fiji^41^. Intensity staining was also calculated using Fiji. Myotube width and intensity values were normalized to the mean value of myotube width and mean intensity of pCLX-pTF-eGFP cells incubated in dox, respectively. Unpaired two sample Mann-Whitney tests were used to compare normalized values using R. P-values were adjusted for multiple comparisons using the Bonferroni method in R.

### Confirmation of doxycycline induced ΔN-p73 protein expression in C2C12 cells

C2C12 cells of wildtype and each mutant genotype were incubated in growth medium plus or minus dox for 48 hours. Cells were then lysed with RIPA buffer (Abcam ab156034) containing Pierce protease inhibitor tablets (ThermoFisher). One hundred micrograms of protein were loaded per well in 4-12% Bis-Tris gels (ThermoFisher) and separated by SDS-PAGE using MOPS buffer (Thermo Fisher). Proteins were transferred to 0.45 um PVDF membranes (ThermoFisher) using a mini blot module (ThermoFisher) and blocked in 5% milk in PBST for one hour. Blots were incubated in primary antibody and 5% milk in PBST overnight at 4C. The primary antibodies used were anti-β-actin (ThermoFisher #MA5-15739) used at a 1:10,000 dilution and anti- ΔN-p73 (ThermoFisher #MA5-16183) used at 1.5 µg/mL. Cells were washed and incubated in secondary antibody for one hour at room temperature. The secondary antibody used was goat anti-mouse-HRP (Thermo Fisher # 31430) used at a 1:20,000 for anti-β-actin blots and 1:5000 dilution for anti-ΔN-p73 blots. Blots were developed using SuperSignal West Pico PLUS Chemiluminescent Substrate (Thermo Fisher#34578) for five minutes before visualization using x-ray film (GeneMate). Western blot images were processed using Fiji.

### Zebrafish modeling of *TP73* loss of function

#### Ethics Statement

All zebrafish experiments were performed in accordance of guidelines from the University of Utah Institutional Animal Care and Use Committee (IACUC), regulated under federal law (the Animal Welfare Act and Public Health Services Regulation Act) by the U.S. Department of Agriculture (USDA) and the Office of Laboratory Animal Welfare at the NIH, and accredited by the Association for Assessment and Accreditation of Laboratory Care International (AAALAC).

#### Fish stocks and animal husbandry

Adult fish were bred according to standard methods. Embryos were raised at 28.5°C in E3 embryo medium with methylene blue and embryos beyond 24 hpf were treated with phenylthiourea (PTU) to prevent pigment formation. Animals of either sex were used for experiments. For *in situ* staining and immunofluorescence, embryos were fixed in 4% paraformaldehyde (PFA) in 1X phosphate buffered saline (1x PBS) overnight (O/N) at 4°C, washed briefly in 1x PBS with 0.1% Tween-20, dehydrated stepwise in methanol (MeOH) (25%, 50%, 75%, 100%), and stored in 100% MeOH at −20°C until use.

#### Immunofluorescence

Immunofluorescence was performed as previously described^42-44^. Antibodies used were: chicken anti-GFP (1:1000 dilution, Aves, GFP-1020), and goat anti-chicken Alexa 488 (1:400 dilution, Invitrogen, A11039). Additionally, nuclei were stained with 4,6-Diamidino-2-Phenylindole (DAPI, diluted 1:1000). Prior to staining, embryos were fixed, dehydrated, and stored as described above. For immunofluorescence, 72 hpf embryos were rehydrated from 100% MeOH to 1x PBST stepwise (MeOH: 100%, 75%, 50%, 25%, 1x PBST), and permeabilized with Proteinase K (1:1000,10 µg/ml) for 1 hour, washed in blocking solution (1x PBST, 1% dimethyl sulfoxide (DMSO), 2% bovine serum albumin (BSA), 5% Normal goat serum (NGS)) for 2 hours, and incubated with primary antibodies diluted in staining solution (1x PBST, 1% DMSO, 2% NGS) overnight at 4°C. The following day embryos were washed several times in 1x PBST. Secondary antibody plus DAPI were diluted in staining solution, and incubated overnight at 4°C. After overnight incubation, embryos were washed several times in 1x PBST and cleared in 25% glycerol in 1x PBS solution for 30 minutes, followed by 50% glycerol overnight at 4°C. The following day embryos were washed in 80% glycerol for 30 minutes and mounted for imaging. For spinal motor neuron (SMN) number quantification, embryos were mounted with their dorsal surface facing toward the coverslip. For axonal outgrowth quantification, the lateral surface of the embryos faced toward the coverslip.

#### TUNEL staining

Terminal deoxynucleotidyl transferase (TdT) dUTP nick-end labeling (TUNEL) was performed on whole-mount larvae (ApopTag Rhodamine *In situ* Apoptosis Detection Kit; Millipore). Embryos were fixed, dehydrated, and permeabilized as described for immunofluorescence. Briefly, embryos were incubated in equilibration buffer (Millipore) for 1 hour, followed by overnight incubation in TdT enzyme (Millipore) at 37°C. The following day, end-labeling was terminated with incubations in stop/wash buffer (Millipore), followed by several washes in 1x PBST. Next, embryos were incubated in working strength sheep anti-digoxigenin rhodamine (Millipore) for 1 hour at room temperature. Labeling was stopped by several washes in 1x PBST and embryos were cleared and mounted for imaging as described above.

#### In situ hybridization

Anti-sense digoxigenin-labeled RNA probe was prepared for *tp73* (ENSDARG00000017953). To generate probe, cDNA from 3 dpf larvae was amplified with the reverse primer containing a T7 RNA polymerase priming site (5’-CCAAGCTTCTAATACGACTCACTATAGGGAGA-3’). The *tp73* gene specific primers used were: forward: 5’-AGCTCAGAGACCACCTCCAA-3’, reverse: 5’- TTCCTCCAACACAACTGCTG-3’. Following PCR amplification, the product was checked for the correct band size by gel electrophoreses, and Sanger sequenced. RNA digoxigenin-labeled probe was synthesized by *in vitro* transcription using Dig RNA labeling kit (Sigma-Aldrich). For *in situ* hybridization, 72 hpf embryos were fixed, dehydrated, and permeabilized as described above. After permeabilization, embryos washed briefly in Hybridization minus (Hyb-) solution (50% deionized formamide (Amresco), 0.1X saline-sodium citrate (SSC), 0.1% Tween 20 (Amresco)) for 5 minutes at 65°C, followed by a 1 hour prehybridization in Hybridization plus (Hyb+) solution (Hyb+ solution contains Hyb- solution plus: 5 mg/ml torula RNA (Sigma-Aldrich), 50 µg/ml heparin (Sigma-Aldrich)) at 65°C. Embryos where then incubated overnight at 65°C with Dig labeled RNA probe diluted in Hyb+ buffer. The following day, embryos were washed at 65°C in 50% formamide in 2XSSCT twice for 30 minutes each, followed by one 2XSSCT wash for 15 minutes, and finally two 0.2XSSCT washes for 30 minutes each. Next embryos were washed in blocking solution (100mM maleic acid, 150 mM NaCl pH 7.5, 2% blocking reagent (Roche)) for 1 hour at room temperature, followed by incubation with anti-digoxigenin alkaline phosphatase (AP) (Roche, 1:5000) overnight at 4°C. The following day, embryos were developed with the addition of BM purple AP substrate (Sigma-Aldrich). *In situ* development was stopped with several washes in 1x PBST and cleared as previously described.

#### In situ hybridization combined with immunohistochemistry

*In situ* hybridization was performed as described above. Embryos were then cryoprotected by washing briefly in 5% sucrose (in 1x PBS) followed by overnight incubation in 30% sucrose at 4°C. Samples were then embedded in optimal cutting temperature compound (OCT, Tissue-TeK, 4583), and stored at −80°C until use. Sections were cut at 20 µm thickness. For immunofluorescence, slides were washed briefly in 1x PBST, and incubated in blocking solution for 30 minutes (1% BSA (Fisher), 0.1% fish skin gelatin (Sigma G7765), 0.1% Tween-20 in 1x PBS). Slides were incubated in primary antibody for one hour at room temperature, followed by several washes in 1x PBST. Next slides were incubated in secondary antibody plus DAPI for one hour at room temperature. Slides were then washed several times in 1x PBST, postfixed in 4% PFA for 10 minutes, and cover slipped using Fluoromount-G (SouthernBiotech).

#### p73 CRISPR mutagenesis

A sgRNA was generated to target the following sequence of exon 4 in zebrafish *tp73*: 5’- CGGCCATCCCTTCCAATACA-3’. sgRNA synthesis was performed as previously described (*23*). Tg(*Hb9:Gal4; UAS:GFP*) embryos were injected with 450 ng/µl *tp73* sgRNA in combination with 800 ng/µl Cas9 protein (IDT). Embryos were collected at 24 hpf for analysis of mutagenesis using high-resolution melt analysis (HRMA) and Sanger sequencing^45^.

Genomic DNA from 24 hpf embryos was prepared as follows: embryos were incubated with 50 µl 50 mM NaOH at 95°C for 25 minutes, followed by neutralization with 5 µl 1M Tris-HCl (pH8.0). PCR reactions were performed in 96-well hard-shell plates (Bio-Rad, Inc.) in 10 µl volume: 4 µl 2.5x LightScanner Master Mix (Biofire, Inc.), 5 pmol of each primer (forward primer: 5-AACCTTCGAGGCCATGTCT-3’; reverse primer: 5’- GTGCTGGACTGCTGGAAAGT-3’), and 2 µl genomic DNA. PCR reactions were covered with 25 µl mineral oil to prevent condensation during HRMA. PCR cycling conditions were: 2 minutes at 95°C, followed by 29 cycles of 30 seconds at 95°C, 30 seconds at 67°C, ending with 74°C for 20 seconds, 25°C for 30 seconds. HRMA was performed on a LightScanner-96 instrument (Biofire, Inc.), from 60°C to 97°C with a temperature transition rate of 0.1°C / s. For sequencing the *tp73* CRISPR targeted region was TA-cloned into a pCR4-TOPO TA (Invitrogen) vector. Four clones per embryo were Sanger sequenced.

#### Transient transgenic zebrafish generation

One-cell stage wild-type embryos were either injected with *mnx*:GFP (18 ng/µl) alone, or in combination (*mnx*:GFP, 50 ng/µl) with *tp73* sgRNA (420 ng/µl). All CRISPR injected embryos were also injected with 800 ng/µl Cas9 protein (IDT). At 72 hpf, embryos were fixed and dehydrated as described above for immunofluorescence and *in situ* staining.

#### Spinal motor neuron (SMN) quantification

To assess the number of SMN, 72 hpf Tg(*Hb9:Gal4; UAS:GFP*) transgenic embryos were stained for GFP, DAPI, and mounted as described. Slides were imaged on a Nikon A1 instrument. A set of 21 slices (step size 5 µm) of the upper third of zebrafish body (starting point below the head) was obtained with identical confocal settings for all embryos (10X objective, 2.5X zoom, 1024 pixels, 6.2 ms / pixel dwell time). Confocal stacks were projected in FIJI and images were composed with Adobe Photoshop (CS6) for quantification of the number of SMN’s per body hemisegment (visible similar blocks of the body coming from mesoderm) using the Photoshop 123 count tool. Representative images were taken on an Olympus FV1000 microscope (30X silicon oil objective, 2048 size, 8 ms / pixel dwell time, 15 slices of 1 µm step size), projected in ImageJ and composed with Adobe Photoshop (CS6) and Illustrator.

#### Spinal Motor Neuron Axonal characterization

To assess axonal outgrowth, transient transgenic 72 hpf immunostained embryos were mounted as described. A set of 21 slices of 5µm step size (50µm above and 50µm below of the area containing the highest number of SMN) was taken in a Nikon A1 microscope using identical settings (10X objective, 2.5X zoom, 1024 size, 6.2 ms / pixel). Confocal stacks of maximal intensity were projected in FIJI and primary and secondary axonal branching length, were measured using Neuron J plugin (scale: 1.9979 pixels/µm; primary axonal length defined as the longest axonal projection from the cell body; for secondary axonal length we average the length of direct branches from the primary axon). Representative images were taken as described above.

#### Apoptosis quantification

To assess apoptosis in SMN, 72 hpf embryos of the above-mentioned stable transgenic line were immunostained and TUNEL labeled, mounted and imaged as described above. Confocal stacks were projected in FIJI and images were composed with Adobe Photoshop (CS6) for quantification of the number of apoptotic SMN out of all SMNs measured in the imaged area. Representative images were taken in a Nikon A1 confocal microscope (20X objective, 2X zoom, 2048 size, 6.9 pixel dwell, 21 slices with 1 µm step size).

#### Statistical Analysis

Statistical analysis was performed using Student’s t-test with a significant p-value set as p<0.05.

**Supplementary Figure 1.**
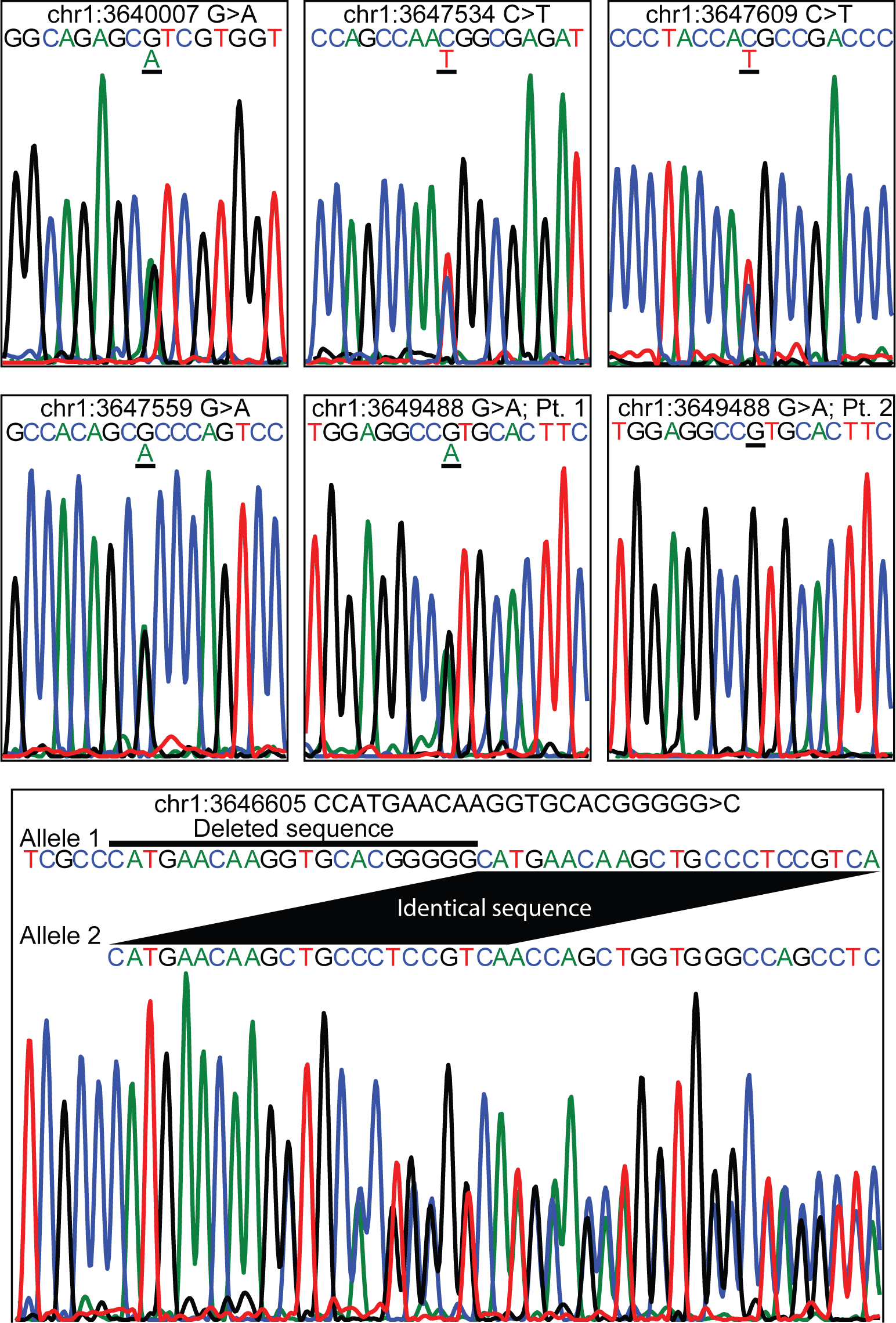
Chromatograms of Sanger validated p73 ALS patient mutations. Sanger sequencing results of the seven *TP73* variants found in the University of Utah ALS patient cohorts. Chr1:3649488 G>A was verified by Sanger sequencing in one of the two patients who were reported to have it by NGS. Pt. stands for patient.

**Supplementary Table 1.**
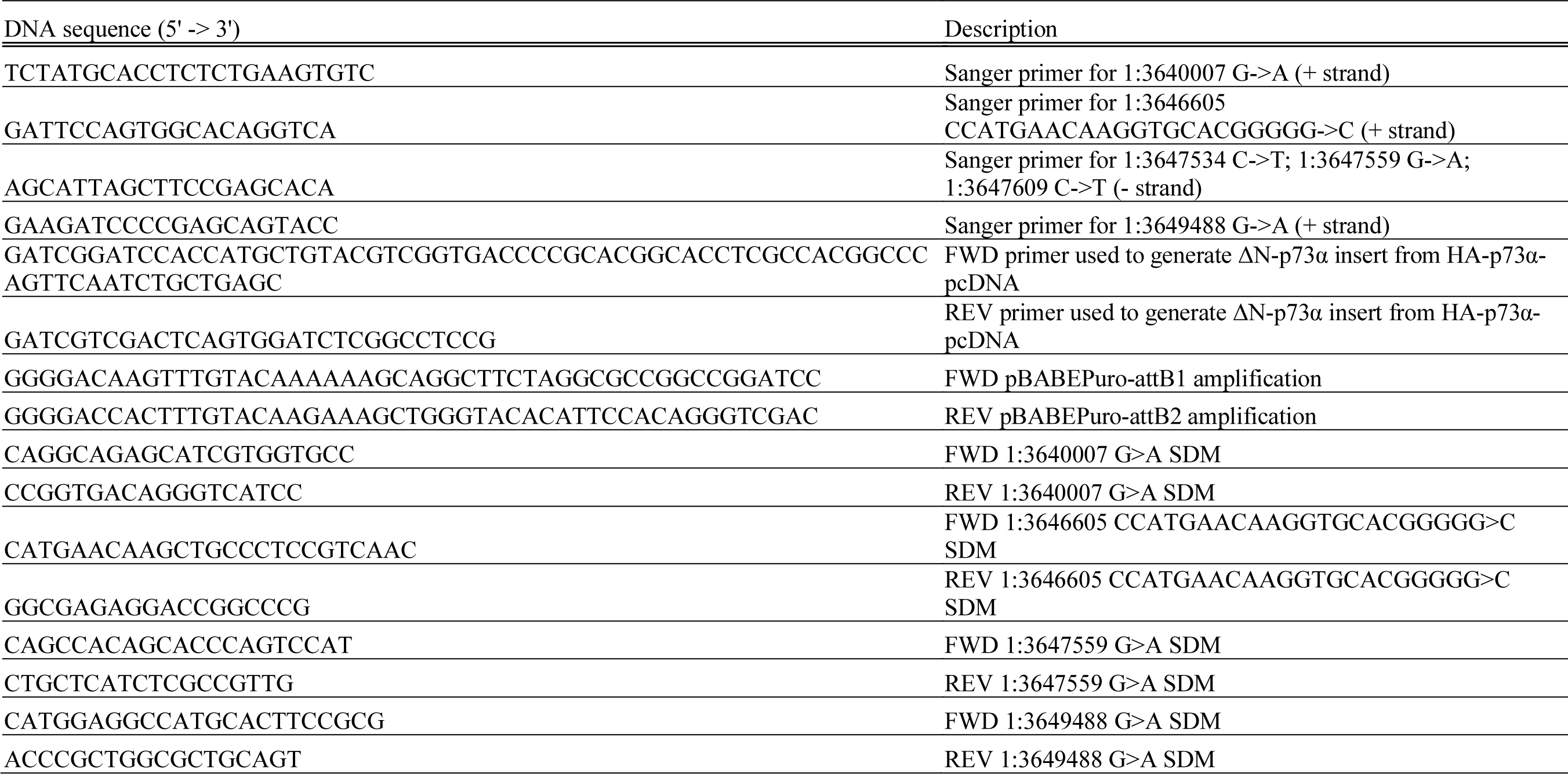
Primers used for site directed mutagenesis and Gateway cloning of WT and mutant ΔN-p73.

